# *Caenorhabditis elegans* AF4/FMR2 family homolog *affl-2* is required for heat shock induced gene expression

**DOI:** 10.1101/817833

**Authors:** Sophie J. Walton, Han Wang, Porfirio Quintero-Cadena, Alex Bateman, Paul W. Sternberg

## Abstract

To mitigate the deleterious effects of temperature increases on cellular organization and proteotoxicity, organisms have developed mechanisms to respond to heat stress. In eukaryotes, HSF1 is the master regulator of the heat shock transcriptional response, but the heat shock response pathway is not yet fully understood. From a forward genetic screen for suppressors of heat shock induced gene expression in *C. elegans*, we identified a new allele of *hsf-1* that alters its DNA-binding domain, and three additional alleles of *sup-45,* a previously uncharacterized genetic locus. We identified *sup-45* as one of the two hitherto unknown *C. elegans* orthologs of the human AF4/FMR2 family proteins, which are involved in regulation of transcriptional elongation rate. We thus renamed *sup-45* as *affl-2* (AF4/FMR2-Like). *affl-2* mutants are egg-laying defective and dumpy, but worms lacking its sole paralog (*affl-1*) appear wild-type. AFFL-2 is a broadly expressed nuclear protein, and nuclear localization of AFFL-2 is necessary for its role in heat shock response. *affl-2* and its paralog are not essential for proper HSF-1 expression and localization after heat shock, which suggests that *affl-2* may function downstream or parallel of *hsf-1*. Our characterization of *affl-2* provides insights into the complex processes of transcriptional elongation and regulating heat shock induced gene expression to protect against heat stress.

## Introduction

Heat is a universal source of stress in nature, which has detrimental effects including disrupting cellular organization and upsetting proteostasis (Morimoto 1998). One way organisms restore homeostasis after heat stress is through rapid transcriptional changes to upregulate genes that assist with combating damaging effects of heat (Morimoto 1998; Hajdu-Cronin *et al.* 2004; Richter *et al.* 2010). In eukaryotes, heat shock induced transcription is initiated when transcription factors known as heat shock factors (HSFs) are activated and bind to heat shock elements (HSEs) in promoters (Morimoto 1998), and HSF1 has been identified as the primary regulator of heat shock induced transcription in eukaryotes (Åkerfelt *et al.* 2010; Richter *et al.* 2010). The current model for transcriptional control of heat shock response by HSF1 is as follows: under normal conditions chaperones sequester HSF1 and upon heat stress the chaperones disassociate with HSF1 so HSF1 is free to upregulate other genes (Richter *et al.* 2010; Voellmy and Boellmann 2007). However, the complete regulatory system that is responsible for the precise transcriptional control of heat shock response is not yet fully understood (Richter *et al.* 2010).

Although regulation of initiation in heat shock induced transcription has been well characterized, it is known that transcription is also tightly regulated at the elongation and termination steps (Kuras and Struhl 1999; Sims *et al.* 2004; Saunders *et al.* 2006; Lenasi and Barboric 2010). For some genes involved in development and stress, RNA Polymerase II becomes paused at the promoter-proximal region, and escape from the paused state becomes a rate limiting process in transcription (Levine 2011; Lin *et al.* 2011; Zhou *et al.* 2012; Luo *et al.* 2012a) In metazoans, Positive Transcription Elongation Factor beta (P-TEFb), a heterodimeric kinase complex composed of CDK-9 and a Cyclin T1 Partner, causes RNA Polymerase II and associated factors to become phosphorylated to stimulate promoter escape (Levine 2011; Zhou *et al.* 2012; Luo *et al.* 2012b). When performing this function, P-TEFb participates in the super elongation complex (SEC): a multi-subunit complex composed of a P-TEFb, an AF4/FMR family protein (AFF1 or AFF4), a Pol II elongation factor (ELL, ELL2 or ELL3), a partner protein (EAF1 or EAF2), and an ENL family protein (ENL or AF9) (Lin *et al.* 2010; Luo *et al.* 2012b). Although both AFF4 and AFF1 serve as scaffolds in the SEC, they have been found to work in different processes; e.g., AFF4 has been found to be more important for HSP70 transcription and AFF1 has been linked to promoting HIV transcription (Lu *et al.* 2015).

*Caenorhabitis elegans* has been used as a multicellular *in vivo* model to perform genetic analysis of homologs of components of the SEC. The *C. elegans* homolog of the ELL gene family (*ell-1*) and its partner (*eaf-1)* are necessary for embryonic development and heat shock induced transcription (Cai *et al.* 2011). In addition, *cdk-9, cit-1.1,* and *cit-1.2*, homologs of CDK9 and cyclin T1, are necessary for phosphorylation of Ser2 in RNA polymerase II in the soma (Shim *et al.* 2002; Bowman *et al.* 2013). However, a *C. elegans* version of the AF4/FMR2 family proteins, which serve as the scaffolds for the SEC, has not yet been identified.

In a genetic screen for suppressors of heat shock induced gene expression, we identified a new reduction of function *hsf-1* allele and cloned a *C. elegans* AF4/FMR2 homolog, *affl-2* (AF4/FMR2-Like). *affl-2* encodes a previously uncharacterized protein that is predicted to have two nuclear localization signals and a collection of disordered residues at its N terminus. Indeed, AFFL-2 is a broadly expressed nuclear protein, and its nuclear localization is necessary for its role in heat shock response. In addition to being defective in heat shock response, *affl-2* mutants are dumpy, egg-laying defective, and some have protruding intestines from their vulvas. *affl-2* is not necessary for the proper localization and expression of HSF-1 pre or post heat shock, suggesting that *affl-2* may function downstream of *hsf-1*. Our identification and characterization of *affl-2* furthers our understanding of heat shock response induced transcriptional regulation in *C. elegans* and validates the power of *C. elegans* for genetic analysis of general transcriptional control.

## Materials and Methods

### *C. elegans* cultivation

*C. elegans* were grown using the methods described in (Brenner 1974). Strains were maintained on NGM agar plates at room temperature (20 °C) and fed OP50, a slow-growing strain of *Escherichia coli*. A list of strains used in this study can be found in Supplementary Experimental Procedures.

### Genomic Editing

We made *affl-1* mutants by inserting the STOP-IN cassette in the 5’ end of the coding sequence of *affl-1* using CRISPR/Cas9 with a co-conversion marker (Wang *et al.* 2018). We injected N2 worms to create *affl-1 (sy1202)* single mutants, and we injected *affl-2(sy975)* worms to create *affl-2(sy975) affl-1(sy1220)* double mutants*. sy1202* and *sy1220* had the same molecular change in the *affl-1* gene. The sequence change is a 43bp insertion with 3-frame stop codon and is near the 5’ end of the gene *Y55B1BR.1/affl-1:*

5’ flanking seq: CCGTACCCGTAGAATGCTTGAAGAAATGGCCGGCC Insertion: GGGAAGTTTGTCCAGAGCAGAGGTGACTAAGTGATAA 3’ flanking seq: TCGTGGGAACTAAACCATTGAGCCAGCTTCCTCGAAG

Primers for genotyping and sequencing can be found in the Supplementary Experimental Procedures.

### Generation of Transgenic Lines

Methods to generate transgenic animals were adapted from Mello and Fire (1995). The *affl-2* driver strain was constructed by injecting 25 ng/µl of pSJW003 along with 40 ng/µl of P*unc-122::rfp* and 35 ng/µl of 1 kb DNA ladder (NEB) into the GFP effector strain *syIs300*, and then outcrossing the resulting worms to add the GFP::H2B effector strain *syIs407* in place of *syIs300* (Wang *et al.* 2017). All *affl-2* rescue variants were constructed by injecting 10 ng/µL of the plasmid containing the rescue construct along with 80 ng/µl of pBluescript and 10 ng/µL of the co-injection marker plasmid KP1368(P*myo-2::nls::mCherry*) into the strain PS8082 (*syIs231 II; affl-2 (sy975) III*). A list of transgenic lines and plasmids used can be found in the Supplementary Experimental Procedures.

### *lin-3c* overexpression assays

We used pumping quiescence as a readout for expression of heat shock driven *lin-3c*, for cessation of pumping is characteristic of *lin-3c* overexpression induced quiescence. We adapted our *lin-3c* overexpression assay from Van Buskirk and Sternberg (2007). 15-30 L4 animals were picked onto NGM agar plates that were seeded 48-72 hours prior. 16-20 hours later, plates with adult animals were parafilmed and placed in a 33 °C water bath for 15 minutes. We used a 15-minute heat shock rather than a 30-minute heat shock because we wanted to be able to detect weaker suppressors. Plates were then left in 20 °C with their lids open to recover for 3 hours before scoring for pumping quiescence. By this time, all worms would have recovered from the mild heat shock and would thus exhibit only *lin-3c* overexpression dependent quiescence. Pumping quiescence was scored using a stereomicroscope on 25-50x magnification, and quiescence was defined as a cessation of movement of the pharyngeal bulb.

### Isolation of suppressors

#### EMS Mutagenesis

Mutagenesis was performed on about 500 late L4 hermaphrodites (PS7244) as described by Brenner (1974). In particular, worms were incubated in a solution of 4 mL M9 with 20 μL EMS (Sigma) for 4 hours. At the end of the 4 hours, we washed the worms three times each with 3 mL of M9 to dilute the EMS. We then plated the mutagenized worms on a seeded NGM agar plate outside the OP50 lawn and left them to recover for at least 30 minutes before plating the P0 worms.

To screen a synchronized F2 population, we treated the F1 adults with alkaline hypochlorite (bleach) treatment to isolate the eggs of the F2 generation (Protocol B from Porta-de-la-Riva *et al.* 2012). After bleaching treatment, we immediately plated the F2 generation eggs. These steps ensured that all of our F2 animals reached adulthood at roughly the same time.

We performed the *lin-3c* overexpression assay as described above on adult F2 worms. We isolated worms who did not exhibit pumping quiescence onto separate plates, and we screened their progeny (F3 generation) to ensure that the phenotype was stable. Mutants isolated from different P_0_ plates were deemed independent.

#### Complementation Testing with *hsf-1* and *affl-2* Mutants

To identify *hsf-1* and *affl-2* mutants, we performed complementation testing with *hsf-1(sy441), affl-2(sy509)*, and *affl-2(sy978)* mutants. Note that *affl-2* was originally named *sup-45. hsf-1(sy441), affl-2(sy509),* or *affl-2(sy978)* hermaphrodites were crossed with *syIs197* males. We crossed the resultant male cross progeny into suppressors, and we performed the *lin-*3c overexpression assay on F1 cross progeny of the cross to assay complementation.

#### SNP Mapping

We used the polymorphic Hawaiian strain CB4856 to perform SNP mapping of our suppressor loci (Doitsidou *et al.* 2010). Our SNP mapping strain was PS7421, which we created by outcrossing PS7244 ten times to CB4856. We followed the Hobert’s Lab protocol for worm genomic DNA for SNP Mapping (pooled samples) (Oliver Hobert Lab) to prepare genomic DNA from the progeny of 50 suppressors and 50 non suppressors crossed with PS7421. For identifying *Y55B1BR.2* we sequenced PS75971 (*syIs231 II; affl-2(sy978) III*) crossed into PS7421 and we sequenced PS8082 (*syIs231 II; affl-2 (sy975) III*) in a N2 background.We analyzed our sequencing results using MiModD mapping software to identify putative mutations (Maier *et al.* 2014).

### Protein Sequence Analysis

Y55B1BR.2 similarity to AF4 was found in the first iteration of a JackHMMER iterative search on the EMBL-EBI webserver (Potter *et al.* 2018). Further rounds of searching revealed homologues in a wide range of eukaryotic species. The search was performed against Reference Proteomes using the phmmer algorithm with the following settings:

#### HMMER Options

-E 1 --domE 1 --incE 0.01 --incdomE 0.03 --mx BLOSUM62 --pextend 0.4 --popen 0.02 --seqdb uniprotrefprot

We used the MUSCLE alignment tool with the default settings to create multiple sequence alignments (Edgar 2004). To determine the location of the predicted NLSs we used the cNLS mapper with a cut-off score of 0.5 and the option to search for bipartite NLSs with a long linker within terminal 60-amino acid regions (Kosugi *et al.* 2008, 2009a; b). We checked that these predictions were consistent with PSORT and NucPred. We used IUPred2A long disorder (Dosztányi *et al.* 2005; Dosztanyi *et al.* 2018) to predict disordered regions of AFFL-2, AFFL-1, AFF1, and AFF4. We used ANCHOR2 software to predict the presence of the disordered protein binding regions of AFFL-2, AFFL-1, AFF1, and AFF4 (Dosztányi *et al.* 2009; Dosztanyi *et al.* 2018). The alignment of the C-terminal homology domain (CHD) was generated using the MUSCLE alignment tool on a selected set of 16 eukaryotic homologues identified using Jackhmmer (Edgar 2004).

### Molecular Biology

We used Roche Taq for genotyping PCR products < 1 kb, and we used NEB Phusion High Fidelity or NEB High Fidelity Q5 for PCR of all cloning inserts and genotyping PCR products that were > 1kb. NEB T4 Ligase was used to construct pSJW003. The NEBuilder HiFi DNA Assembly Master Mix was used for Gibson Assembly of pSJW005, pSJW035, pSJW036, pSJW040, and pSJW041. We used DH5α competent cells for all transformation, and we plated all transformed cells on carbenicillin LB plates. A detailed list of oligos and plasmids can be found in the Supplementary Experimental Procedures.

### Microscopy

For images of plate phenotypes, one day old adult worms were filmed on plates seeded within 24 hours after the L4 larvae stage. We used a Wild Makroskop M420 dissecting scope at 32x or 40x magnification.

Images of *affl-2* expression and localization of HSF-1::GFP and AFFL-2::GFP were taken using a Plan Apochromat 10x or 63x/1.4 Oil DIC objective in a Zeiss Imager Z2 microscope with an Axiocam 506 Mono camera using ZEN Blue 2.3 software. Young adults were used for imaging, because their lower fat content made it easier to detect fluorescent proteins. Worms were anesthetized in 3 mM levamisole and mounted on 2 % agarose pads. Z-stacks were taken at 63x and maximum-intensity projections were generated using Fiji/ImageJ software (Schindelin *et al.* 2012). Z-stacks were used for images of HSF-1::GFP localization, and when indicated for AFFL-2::GFP localization.

Our method to image HSF-1::GFP was adapted from Morton and Lamitina (2013). To image HSF-1::GFP after heat shock, worms were mounted on slides and incubated in a preheated PCR machine at 35 °C for five minutes. The lid of the PCR machine was left open, for it did not reach 35 °C. Instead, slides were placed in a foil packet on top of the PCR wells. We found that a five-minute heat shock was long enough to cause worms to form granules consistently and longer incubation caused the plates to dry out. Worms were imaged immediately after heat shock. Non heat shock controls were left on the benchtop (20 °C) for five minutes prior to imaging. In our image analysis, we selected nuclei manually, and chosen nuclei were segmented automatically. We detected granules in chosen nuclei automatically using the blob detector function from scikit-image (Walt *et al.* 2014). We used the mean intensity of the segmented nuclei and granules after background subtraction to determine HSF-1::GFP nuclear intensity.

To image AFFL-2::GFP after heat shock, worms were heat shocked following the same protocol as we used for the *lin-3c* overexpression assay. We choose to use this protocol rather than the one adapted from (Morton and Lamitina 2013) because we wanted to recreate the conditions of the worms were in during the genetic screen. After the heat shock worms were immediately mounted onto slides and imaged.

### Data Analysis

Unless otherwise specified, data analysis was carried out using Python 3.7 with standard scientific libraries (Jones *et al.* 2001) and version 0.041 of the bebi103 package (Bois 2018).

#### Determining pumping quiesence frequency

We used a Bayesian modeling framework to estimate the frequency of worms pumping after heat shock in the *lin-3c* overexpression assay (Bois 2018)

Our prior distribution for *ϕ* was:

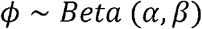

Where, *α*, *β* are parameters for the Beta distribution that we specified. Our likelihood for *n*, the number of worms exhibiting pumping quiesence was

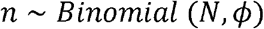

where *N* is the number of worms in each experiment, *n* is the number pumping, and *ϕ* is the probability of pumping. We used Stan to sample from the posterior distribution and find confience intervals for the parameter estimates (Stan Development Team, 2018).

#### Quantification of HSF-1 Expression and Nuclear Granule Formation

We used non-parametric bootstrapping to estimate the mean of HSF-1 nuclear intensity before heat shock, HSF-1 granule intensity after heat shock, and estimate of the mean of the number of granules per nuclei formed in each strain. We used the permutation test to compute all reported p-values, for we did not have a good parametric model to explain the data generating processes.

## Results

### A genetic screen of pumping quiescence suppression yields new alleles involved in heat shock transcription response

To identify genes involved in heat shock response, we conducted a forward genetic screen to search for suppressors of pumping quiescence caused through P*hsp-16.41* (promoter of *hsp-16.41*) driven *lin-3* (Fig 1A). When overexpressed in adult animals, *lin-*3, encoding the *C. elegans* ortholog of the epidermal growth factor (EGF), causes a reversible state of quiescence that is characterized by cessation of feeding, locomotion, defecation, and decreased responsiveness (Van Buskirk and Sternberg 2007). *hsp-16.41*’s expression is induced by heat shock; therefore, placing *lin-3* under the control of P*hsp-16.41* gives us temporal control of quiescence. We screened for animals which are defective in downstream steps of quiescence or do not express the transgene properly by searching for pumping animals after heat shock (Fig 1A). We expected that some suppressors would be defective in heat shock induced transcription of the *lin-3* transgene. Indeed, we found that one of the strong suppressors, *sy1198*, is recessive and was mapped to the chromosome I (data not shown). The gene *hsf-1* is also located on chromosome I, and it was identified from a similar screen using a transgene with a gain-of-function *goa-1* driven by the promoter of *hsp-16.2,* another heat shock response gene similar to *hsp-16.41* (Hadju-Cronin et. al, 2004). We found that *sy1198* and *hsf-1(sy441)* failed to complement for suppression of quiescence, which indicated that *sy1198* is an allele of *hsf-1.* By Sanger sequencing, we found *sy1198* is a T to A mutation in the *hsf-1* gene, which leads to a Leucine to Glutamine substitution (L93Q) in the predicted DNA binding domain of HSF-1 protein (Table 1, Fig 1b; Hadju-Cronin *et al.* 2004). As null mutants of *hsf-1* are lethal (Li *et al.* 2016), *sy1198* is likely to be a weak loss-of-function allele.

**Figure 1.**
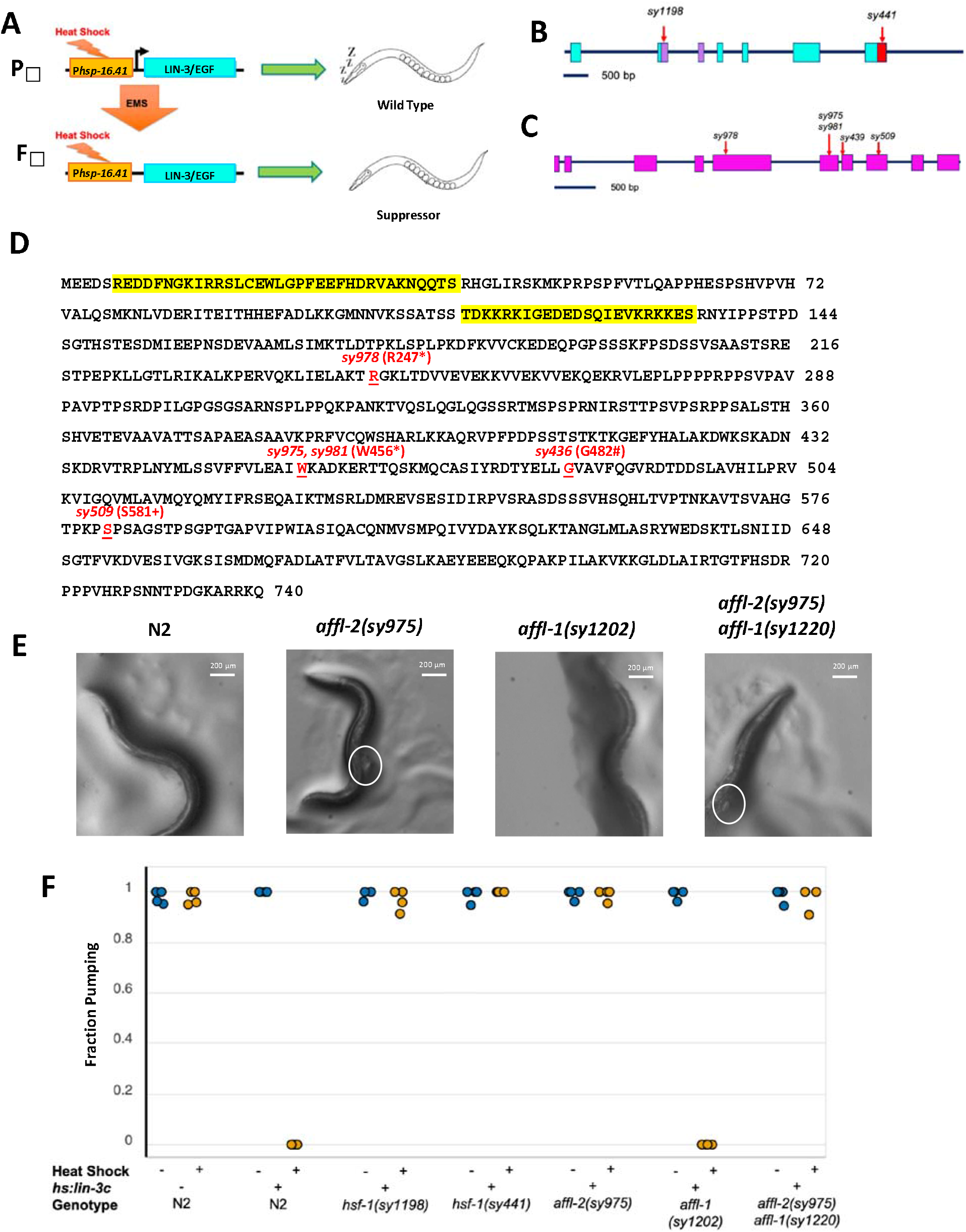
*hsf-1* and *affl-2* were discovered in a screen for suppressors of heat shock induced *lin-3c* overexpression. (A) Outline of screening process. The F2 progeny of mutagenized Phsp-16.41*:lin3-c* animals were screened for suppression of *lin-3c* overexpression induced pumping quiescence. Gene diagrams of *hsf-1* (B) and *affl-2* (C). Blocks are exons, lines are introns, and red arrows indicate position of molecular changes for alleles. Sequences for alleles can be found in table 1. (D) Annotated AFFL-2 sequence. Highlighted text in yellow corresponds to predicted NLSs, and disrupted residue due to allele change is underlined in red text. * indicates stop; # indicates splice site disruption prior to residue; + indicates first disrupted residue of missense mutation and frameshift caused by *sy509.* (E) Images of wild type (N2), *affl-2* mutant*, affl-1* mutant, and *affl-2 affl-1* double mutants. Protruding intestines from vulvas are circled. (F) Fraction of worms pumping before or after heat shock to induce *hsp-16.41: lin-3c* expression. Data are a proxy for gene expression (pumping quiescence indicates *hsp-16.41* expression). Each point represents a different trial.

**Table 1:**
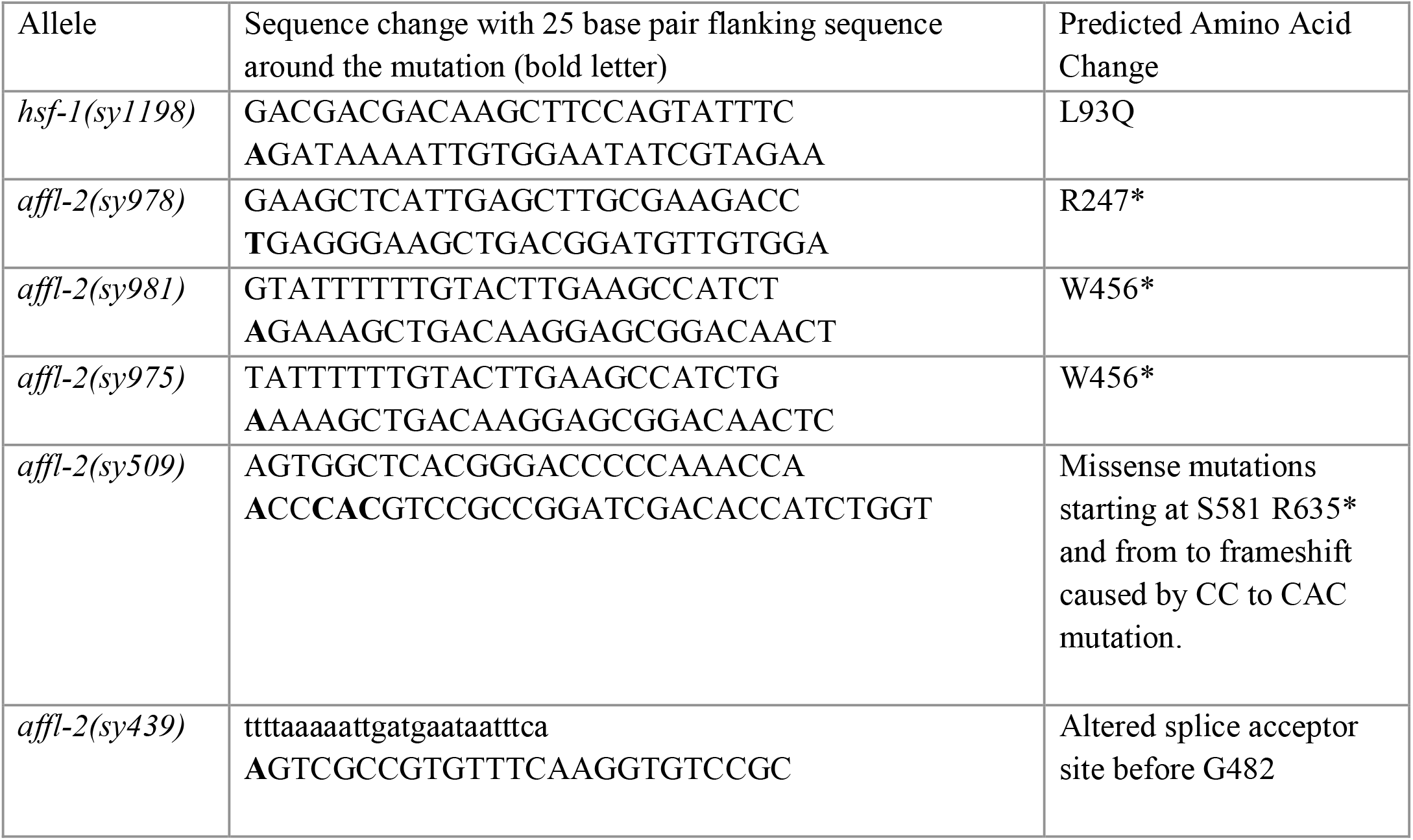
DNA Sequence Changes of Alleles. Alleles *sy439* and *sy509* were both identified in Hadju-Cronin et. al (2004), where *affl-2* was named *sup-45.* All other alleles were identified in this screen. Bold letters indicate modified bp(s) in that allele. Uppercase represents exons and lowercase represents introns.

### AFFL-1 and AFFL-2 are homologs of members of the AF4/FMR2 family

We used Hawaiian SNP mapping and whole genome sequencing to find candidates for causal mutations of the suppression of quiescence. We noticed that a group of suppressors which failed to complement one another had mutations in the uncharacterized gene *Y55B1BR.2,* and their mapping position (Fig S1) was similar to that of *sup-45*. Hadju-Cronin *et al.* identified *sup-45* mutants that are defective in heat shock induced transgene expression, but they were unable to clone the gene. We found that our *Y55B1BR.2* mutants failed to complement *sup-45(sy509)*, which confirmed that the suppressors are alleles of *sup-45.* We determined the molecular changes of our three *sup-45* alleles and two previously described *sup-45* using Sanger sequencing, and indeed all had mutations in *Y55B1BR.2* (Table 1). For the rest of our experiments, we used *sy975,* which contains a nonsense mutation at residue 456 (Fig 1C-D, Table 1).

We found that Y55B1BR.2 is predicted to have two nuclear localization signals (NLS), which suggested that it is a nuclear protein (see Methods). Additionally, Y55B1BR.2 has an adjacent paralog *Y55B1BR.1*, whose encoding gene’s start codon is separated from the stop codon of *Y55B1BR.2* by only 489 nucleotides (“Y55B1BR.2 (gene) - WormBase : Nematode Information Resource). Using JackHMMR (Potter *et al.* 2018), we found that Y55B1BR.2 and Y55B1BR.1 are homologs of AFF4, which is a member of the human AF4/FMR2 family. Based this homology we chose to name *Y55B1BR.1* as *affl-1* (AF4/FMR2 Like) and *Y55B1BR.2* as *affl-2.* The AF4/FMR2 family includes the proteins AFF4 and AFF1, which serve as scaffold proteins for multi-subunit super elongation complexes (SECs) that assist with releasing RNA polymerase from promoter-proximal pausing (He and Zhou 2011; Lu *et al.* 2014; Mück *et al.* 2016. AFF1 and AFF4 both consist of an intrinsically disordered N-terminus that interacts with other members of the SEC and a C-terminal homology domain (CHD) that is conserved among members of the AF4/FMR2 family (Chen and Cramer 2019). AFF4’s binding sites to SEC partners have been studied and are diagramed in Fig 2B. Similarly, AFFL-2 is also predicted to have disordered residues, most of which are in its N-terminus (Fig 2A). Disordered proteins sometimes have protein binding domains that are disordered in isolation but become structured upon binding (Dosztányi *et al.* 2009), and AFFL-2 has three such predicted disordered protein binding regions in its N-terminus (Fig 2A).

**Figure 2.**
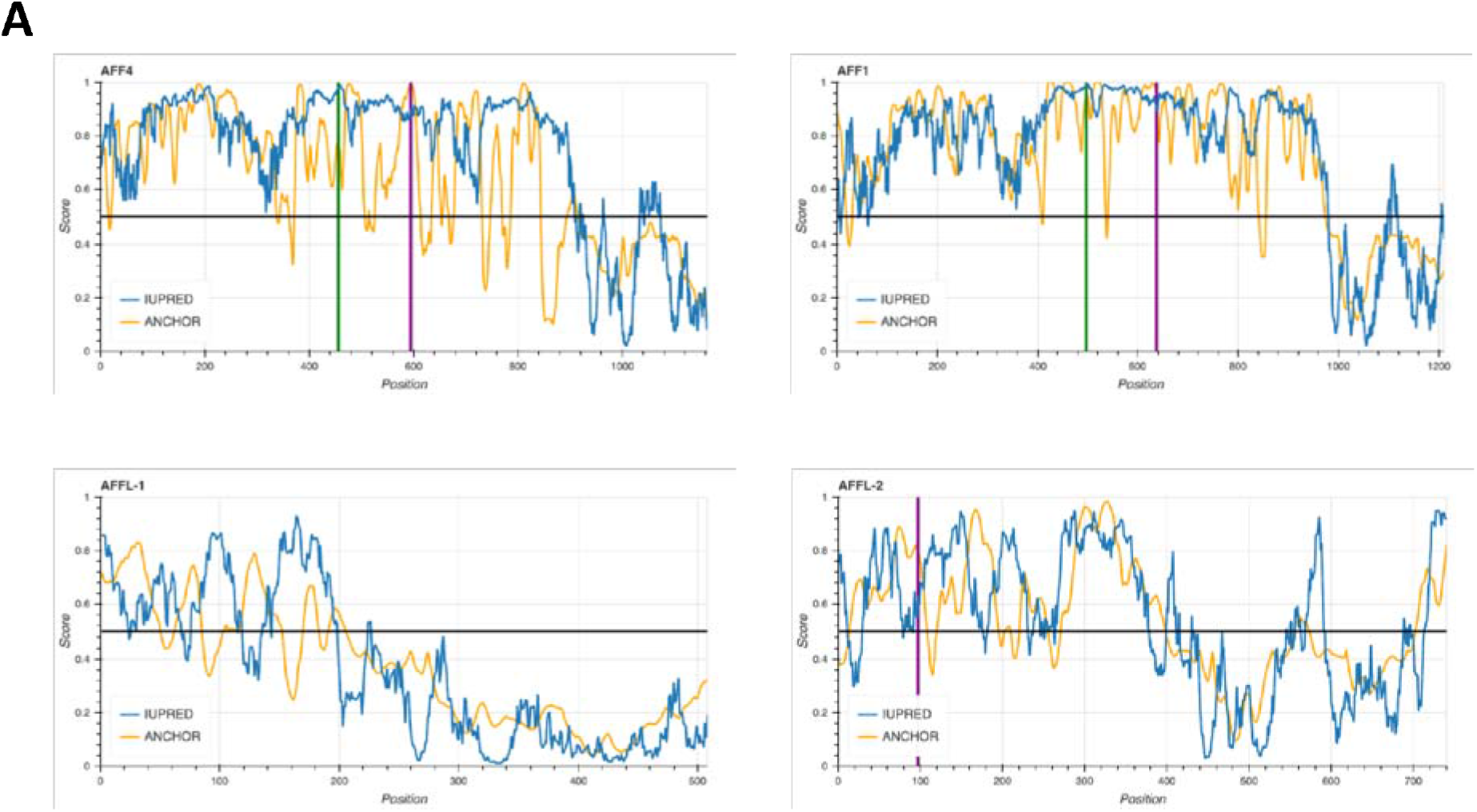
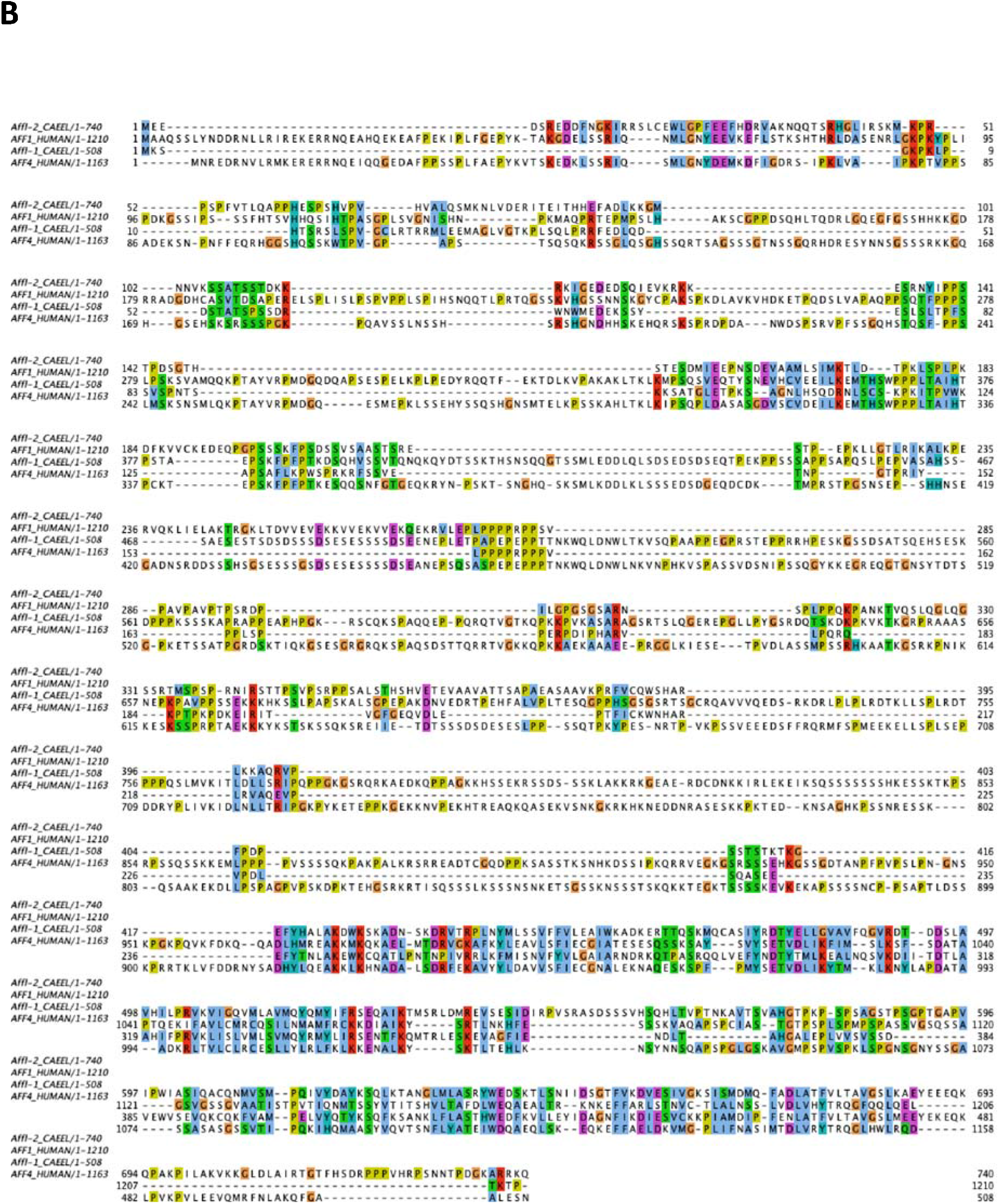
*affl-*2 is a homolog of the AF4/FMR2 family and predicted to be a disordered protein. (A) Plots of ANCHOR and IUPRED score per residue for AFF4, AFF1, AFFL-1, and AFFL-2. IUPRED predicts disordered residues, and higher scores indicate higher confidence that a residue is disordered (Dosztányi *et al.* 2005; Dosztanyi *et al.* 2018). ANCHOR predicts disordered protein binding regions, and higher scores indicate higher confidence that a disordered residue can participate in protein binding (Dosztányi *et al.* 2009).(B) Diagram of *H. sapiens* AFF4 with annotated binding sites (Chou *et al.* 2013; Leach *et al.* 2013; Schulze-Gahmen *et al.* 2013, 2014; Qi *et al.* 2017; Schulze-Gahmen and Hurley 2018; Chen and Cramer 2019). (C) Sequence alignment of AFFL-2, AFFl-1 (AFFL-2’s paralog in *C. elegans*), and AFF1 and AFF4 in *H. sapiens*. The alignment was prepared using Jalview with ClustalX coloring to highlight conservation (Edgar 2004), where conserved residues are colored as follows: hydrophobic (blue), positive charge (red), negative charge (magenta), polar (green), cysteines (pink), glycine (orange), proline (yellow) and aromatic (cyan). The UniProtKB accession identifiers for each sequence are listed here: Affl-2_CAEEL, Q95XW7; Affl-1_CAEEL, Q95XW6; AFF1_HUMAN, P51825; AFF4_HUMAN, Q9UHB7.

Multiple Sequence Alignment of *H. sapiens* AFF1and AFF4 along with *C. elegans* AFFL-2 and AFFL-1 reveals that most of the similarity between the four proteins is in the conserved C-terminal Homology Domain (CHD) of the AF4/FMR2 family members (Fig 2C, Fig S2).The CHD of AFF4 has been shown to bind nucleic acids, form homodimers, and form heterodimers with the AFF1-CHD *in vitro* (Chen and Cramer 2019). Interestingly, our sequence searches identified previously unknown homologues of AF4/FMR2 proteins in *Arabidopsis thaliana*, *Dictostelium discoideum*, *Acanthamoeba castellani* and sporadic yeast species. Gopalan and colleagues previously identified a homologue within *Schizosaccharomyces pombe* (Gopalan *et al.* 2018).

### *affl-2* mutants, but not *affl-1* mutants, are Egl, Dpy, and deficient in heat shock response

*affl-2* mutants have been found to be Egl (EGg Laying defect), Dpy (Dumpy) and deficient in heat shock induced transcription (Hajdu-Cronin *et al.* 2004). We also noticed that some *affl-2* mutants have their intestines protruding from their vulvas (Fig 1E). We did not observe such hernias in either *hsf-1(sy441)* or *hsf-1(sy1198)* worms.

We did not find any *affl-1—*the paralog of *affl-2—* mutants in our screen, so we made a null mutant of *affl-1* to see if it is also necessary for heat shock induced gene expression. *affl-1* mutants appear wild-type and do not have any of the morphological phenotypes characteristic of *affl-2* mutants (Fig 1E). We also made *affl-2(sy975) affl-1(sy1220)* double-mutant animals, which we found are Dpy, Egl, and have herniated intestines. We did not notice any obvious defects in the double mutants that are not present in *affl-2(sy975)* single mutants (Fig 1E).

*affl-2* has been identified in two independent forward genetic screens using different transgenes: one driven by *Phsp-16.2* (where it was called *sup-45)* and the other by *Phsp-16.41.* It has already been demonstrated that *hsp-16.2* transcription is eliminated in *hsf-1* and *affl-2* mutants worms using qPCR (Hajdu-Cronin *et al.* 2004). Since *hsp-16.2* and *hsp-16.41* share the same regulatory sequence and are both induced by heat shock (Jones *et al.* 1986), we believe that *hsp-16.41* transcription is also likely eliminated in *hsf-1* and *affl-2* mutants. Thus, we decided to use heat shock inducible pumping quiescence, due to expression of *Phsp-16.41*, driven *lin-3* as a readout for *hsp-16.41* expression. We estimated *ϕ*, the probability of a given worm pumping after heat shock, to see whether the *hsp-16.41* promoter is active under heat shock conditions in *affl-2, hsf-1, affl-1,* and *affl-2 affl-1* mutants (Fig 1F). We also ensured that all mutants pump at wild type levels prior to heat shock (Fig 1F). The estimates of for *hsp-16.41:lin-3c* and *affl-1(sy1202); hsp-16.41:lin-3c* were both 0, which indicates that there are no defects in heat shock induced *hsp-16.41* expression. The estimates of for wild type, P*hsp-16.41:lin-3c; affl-2(sy975),* P*hsp-16.41:lin-3c; hsf-1(sy441),* P*hsp-16.41:lin-3c; hsf-1(sy1198),* P*hsp-16.41: affl-2(sy975) affl-1(sy1220)* were all close to one, which indicates that these mutants were not expressing the *lin-*3c transgene after heat shock. These results confirmed that *hsf-1* and *affl-2,* but not *affl-1,* are necessary for heat shock induced *hsp-16.41* expression.

### AFFL-2 is a broadly expressed nuclear protein

We cloned the first 3 kb of sequence upstream from *affl-2’s* start codon as *affl*-2’s 5’ regulatory region and promoter. We used this sequence to create a cGAL driver (Wang et al. 2017), which we crossed with a GFP::H2B effector to create a transcriptional reporter for *affl-2*. GFP was visible in all tissues in worms of all stages, which indicates that *affl-2* is ubiquitously expressed (Fig 3A). To observe the subcellular localization of AFFL-2, we used our cloned *affl-2* promoter to drive AFFL-2 cDNA::GFP. *affl-2(sy975)* mutants with the transgene appear wild-type and are able to express the *hsp-16.41* promoter driven *lin-*3c transgene (Fig 3E). Therefore, we believe that our fusion protein is functional, and our cloned promoter for *affl-2* reflects its endogenous expression pattern. AFFL-2::GFP is exclusively located in the nucleus prior to heat shock, and we do not see any noticeable difference in AFFL-2::GFP localization or intensity between worms before and after heat shock (Fig 3B).

**Figure 3.**
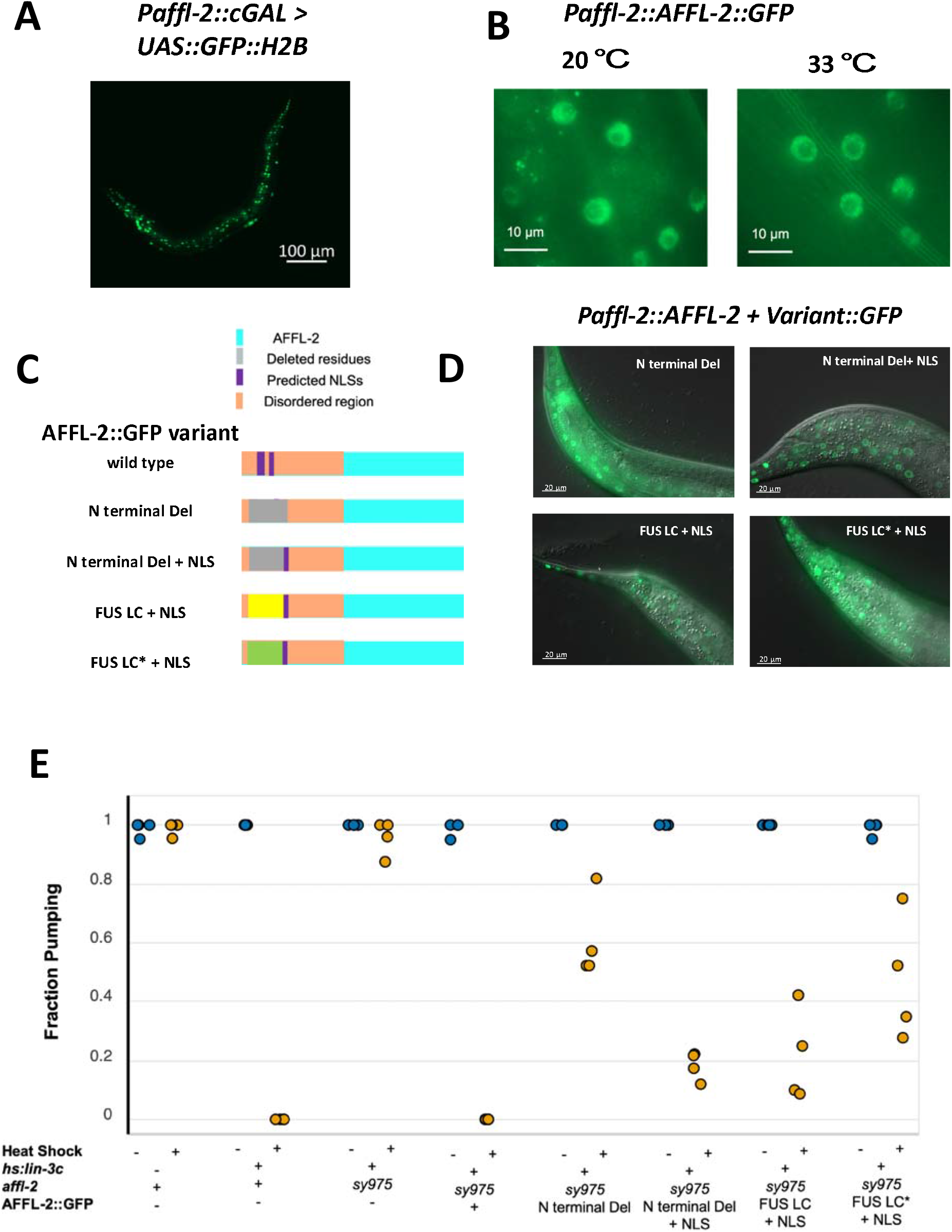
AFFL-2 is a ubiquitously expressed nuclear protein. A) Representative image of a worm expressing *Paffl-2*::cGAL > *15xUAS::GFP::H2B,* which can be seen in the majority of cells, indicating that *affl-2* is ubiquitously expressed. Note that the effector construct is integrated, while the driver is an extrachromosomal array. B) AFFL-2::GFP at 20 °C and 33 °C. Both images are taken of nuclei in the tail of young adults. (C) Diagrams of AFFL-2 variants. Note that diagrams are not to scale, but are just representative of the ordering of various elements. FUS LC* represents the modified FUS LC residues with disordered residues mutated to more ordered ones. All constructs are driven by the *affl-2* promoter (P*affl-2*). (D) Subcellular of localization of AFFL-2 variants. Animals in photos are young wild type adults at room temperature. (E) Fraction of worms pumping before or after heat shock to induce *hsp-16.41: lin-3c* expression. Data are a proxy for gene expression (pumping quiescence indicates *hsp-16.41* expression). Each point represents a different trial.

As shown previously, *affl-1* is not necessary for heat shock induced *hsp-16.41* transcription despite also being a homolog of AFF1 and AFF4 (Fig 1E). Additionally, our Multiple Sequence Alignment suggests that AFFL1 shares little similarity with the first 135 amino acids of AFFL2, and AFFL-1 is not predicted to have any NLS (Fig 2, see Methods). We thus decided to investigate the role of AFFL-2’s N-terminus, which contains its predicted NLSs and the majority of its predicted disordered residues (Fig 1D, 2A) by creating alternative versions of AFFL-2 with modified N termini (Fig 3C). First, we created a modified AFFL-2::GFP in which we removed 129 amino acids from the N terminus of AFFL-2. This modification eliminated both predicted NLSs and the majority of the disordered residues. To test the necessity of the disordered residues independently from the NLS, we created a construct in which we substituted the deleted residues with the SV40 NLS. To test the role of the disordered nature of the domain independently of its sequence, we made another construct that included the artificial NLS and the 212 residue fused in sarcoma low complexity (FUS LC) domain. FUS is one of three RNA binding proteins with LC domains that when fused to DNA binding domains cause a variety of cancers (Arvand and Denny 2001; Guipaud *et al.* 2006; Lessnick and Ladanyi 2012). 84 % of the FUS LC domain consists of glycine, serine, glutamine, and tyrosine, and at high concentrations the domain has been shown to polymerize. Kwon *et. al.* (2013) also demonstrated that the FUS LC domain fused to the GAL4 DNA binding domain can induce transcriptional activation. As a control, we added a similar region but with 10 tyrosines mutated to serines, which was shown to prevent the GAL4 and FUS LC fusion transcriptional activation abilities (Kwon *et al.* 2013).

We found that in a wild-type background, all of the altered AFFL-2 proteins containing an NLS are observed in the nucleus (Fig 3D). The modified AFFL-2 lacking the artificial NLS is primarily located in the nucleus, but we saw that some of the protein is present in the cytoplasm (Fig 3D, Fig S3). To test whether the nuclear localization of this construct is dependent on the presence of wild type AFFL-2, we introduced the modified AFFL-2 with N-terminal deletion in *affl-2(sy975)* animals. The localization of AFFL-2::N-terminal Deletion::GFP was similar in both a wild type and *affl-2(sy975)* background: some animals had it strictly localized to the nucleus while others had it in the cytoplasm as well (Fig S3). We did not expect this modified protein to localize to the nucleus, but this result suggests there is an alternative mechanism for its nuclear import. Introducing the modified AFFL-2 with the deletion, but no other alterations, did not rescue any morphological defects of *affl-2(sy975)* worms (Fig S4). However, the constructs with an artificial NLS rescued the morphological defects of *affl-2(s975)* mutants (Fig S4).

To determine the extent that the constructs rescued the heat shock induced gene expression defects of *affl-2(sy975),* we estimated the probability of worms exhibiting pumping quiescence due to heat shock induced *Phsp-16.41: lin-3c* expression (Fig 3E). The estimate of (the probability of pumping after heat shock) for *affl-2(sy975)* animals with a wild type copy of AFFL-2::GFP was 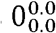 (estimated mean with lower and upper subscripts denoting lower and upper bounds for estimated 95% confidence interval), which demonstrates that the full AFFL-2::GFP construct is functional. The estimates for *ϕ* for the deletion-only construct and the construct with the addition of the modified FUS LC domain were both relatively high 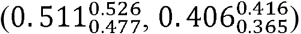, and the estimates for *ϕ* for constructs with only the artificial NLS and the FUS LC were much lower (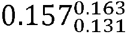 and 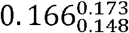, respectively). Even though the modified AFFL-2 with both predicted NLSs removed can be seen in the nucleus, adding back the artificial NLS significantly increased the performance of AFFL-2. This suggests that exclusive restriction of AFFL-2 in the nucleus is important for its function in heat shock response. Adding back the low complexity FUS LC did not further increase the performance of the modified AFFL-2, but the modified (Tyrosine to Serine) FUS LC hindered the performance of AFFL-2. It is believed that FUS increases transcriptional activity by enhancing recruitment of RNA polymerase II, for mutants that bind RNA polymerase II better increase transcription (Kwon *et al.* 2013), and thus this result suggests that the role of AFFL-2 is not to enhance recruitment of RNA polymerase

### *affl-1 and affl-2* do not significantly influence HSF-1 localization and expression

Since *hsf-1* is an essential gene, we could not perform traditional epistasis experiments using the null mutants to determine whether, and if so how, *affl-2* and *hsf-1* genetically interact. Instead, we used an *hsf-1* translational reporter to determine if *affl-2* and/or *affl-2* are necessary for proper localization and expression of HSF-1. *C. elegans* HSF-1 is a ubiquitously expressed nuclear protein, and HSF-1 will aggregate to form nuclear stress granules after heat shock (Morton and Lamitina 2013). The HSF-1 foci do align with marks of active transcription and are dependent on the HSF1 DNA binding domain, but the putative sites of the foci are still unknown (Morton and Lamitina 2013). We quantified the formation of granules after heat shock using the *Phsf-1::*HSF-1::GFP transgene from Morton and Lamitina (2013) (Fig 4). We also quantified the intensity of the granules after heat shock and the intensity of HSF1::GFP prior to heat shock for all genotypes (Fig S5). We found that HSF-1 expression prior to heat shock is similar in all genotypes (Fig S5a), which demonstrates that neither *affl-2* nor *affl-1* are critical for regulating HSF1 expression. Although some differences of means of the number of granules (Fig 4b) and intensity (Fig S5b) of HSF-1::GFP between genotypes are statistically significant (*p* < 0.05), there is still too much overlap between the different distributions of HSF-1::GFP intensity for these differences to fully explain the highly non-overlapping differences between the quiescence phenotype of the different strains. Surprisingly*, affl-1(sy1202)* worms have more granules per nucleus after heat shock compared to wild-type worms (*p* = 0.0016 and the most number of granules compared to the other mutants. *affl-2(sy975)* mutants formed slightly less granules after heat shock (*p* = 0.034) and *affl-2(sy975) affl-1(sy1220)* mutants formed similar numbers of granules per nucleus as wild type (*p* 0.883). These results do not explain why *affl-2(sy975)* worms are unable to express *hsp-16.41,* for some wild type worms had similar HSF1 granule numbers per nuclei and similar HSF-1::GFP levels. Since HSF-1 localization and expression is not significantly disrupted in *affl-2* mutants, we believe that AFFL-2 acts either downstream or parallel to HSF-1 to regulate heat shock induced transcription in *C. elegans*. This suggests a predicted role of *affl-2* in elongation, based on its homology with AF4/FMR2 family members.

**Figure 4.**
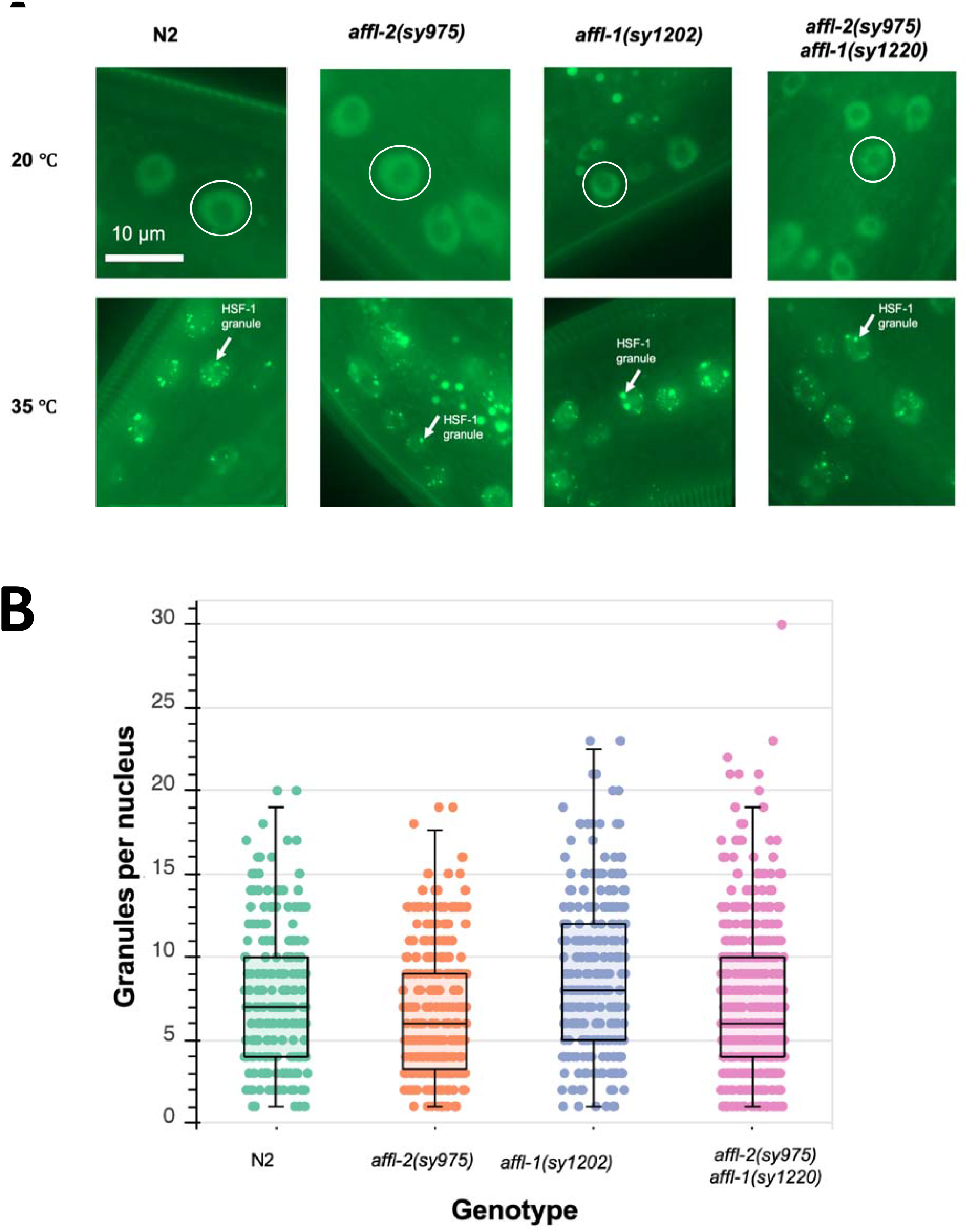
Analysis of the interaction of *hsf-1, affl-2,* and *affl-1.* All genotypes contain *drSi13,* which is the single-copy transgene with P*hsf-1::HSF-1::GFP.* (A) Representative images of hypodermis nuclei in young adults. Subcellular localization of HSF-1::GFP with and without a five-minute heat shock at 35 **°**C. Prior to heat shock, HSF-1::GFP is distributed throughout the nucleus for all genotypes; after heat shock HSF-1::GFP forms nuclear granules. Images of nuclei are from the hypodermis but are representative of HSF-1::GFP localization throughout the entire animal. In top row, representative nuclei are circled. In bottom row, examples of nuclear granules are indicated with arrows. (B) Quantification of HSF-1::GFP granules per nucleus after heat shock for different genotypes. Each point represents one nucleus.

## Discussion

We have cloned and performed a genetic analysis of *affl-2*, a homolog of AF4/FMR2 family members and showed that *affl-2* is necessary for heat shock induced transcription. Through a forward genetic screen for suppressors of heat shock induced *lin-3* overexpression, we identified one new *hsf-1* and three *affl-2* alleles. To our knowledge, this is the first isolated viable *hsf-1* allele with an altered DNA binding domain. We found that *affl-*2 mutants are Dpy, Egl, and have herniated intestines, whereas animals lacking a functional *affl-1*—a homolog of AF4/FMR2 family members and the paralog of *affl-2*— appear wild type. We determined that *affl-2* is a ubiquitously expressed nuclear protein, and proper localization is necessary for its role in heat shock induced transcriptional response.

*affl-2* mutants had been identified in another screen for suppressors of heat shock induced gene expression, and that screen also identified *hsf-1* and *cyl-1* as regulators of heat shock response (Hajdu-Cronin *et al.* 2004). As said above, we found an *hsf-1* mutant, but we did not recover any *cyl-1* mutants. However, Hajdu-Cronin *et al.* (2004) reported that their *cyl-1* mutants did not suppress the effects of a P*hsp-16.41* driven transgene, which indicate that *cyl-1* is not responsible for *hsp-16.41* transcription. Hadju-Cronin’s efforts and ours illustrate the power of simple genetic screens for gene expression in *C. elegans* to find genes responsible for regulation of transcription.

Along with conservation in the CHD, we see a partial conservation of AFF4’s AF9/ENL, ELL-1/2, and P-TEFb binding sites in the AFFL-2 N terminal sequence (Fig 2C). AFFL-2 is predicted computationally to have three candidates for binding sites, but we have not yet verified whether these are real. It is possible that these three sites are sufficient for AFFL-2 to bind to its partners in the SEC, and it is possible that AFFL-2 does not interact with all components of the SEC that human AFF4/AFF1 have been found to bind. Our deletion removed the region of AFFL-2 similar to AFF4’s binding site to P-TEFb, and thus we expected it to be necessary for the AFFL-2’s function. Surprisingly, we found that replacing much of the disordered N-terminus of AFFL-2 with an exogenous NLS restores protein function to about 80% of the wild-type control, even though the modified AFFL-2 with its predicted NLSs at the N-terminus removed still partially localizes to the nucleus. It is possible that the C-terminus of AFFL-2 may contain a weak NLS that cannot be predicted by current software which allows AFFL-2 to partially localize to the nucleus at low levels. Addition of an exogenous NLS could be necessary to increase the concentration of nuclear AFFL-2 to improve its functioning, but does not restore AFFL-2 activity to wild-type levels. Our deleted residues removed only one of the candidate binding sites of AFFL-2, and it is possible that the other binding sites and disordered residues can act redundantly to maintain AFFL-2 activity. However, a more thorough biochemical investigation of AFFL-2 is needed to determine the role of different domains of the protein.

Despite AFFL-1 being an ortholog of AFF4/AFF1 as well, AFFL-1 is not necessary for heat shock induced transcription and *affl-1* mutants appear wild type. AFFL-1 is not predicted to have any Nuclear Localization Signals, which suggests that it may not even be a nuclear protein. Since we do not have a phenotype for *affl-1* mutants we have no way to validate any expression pattern or localization obtained using a fusion protein, for we cannot validate that the fusion protein is functional. It is not clear what AFFL-1’s role is, and if AFFL-1 has a role in transcription or not. We have not fully investigated *affl-1* mutants to see if they have any deficiencies in other processes besides heat shock induced transcription, and *affl-1* could play a redundant role with another gene.

We used translational reporters to examine the roles of *affl-1* and *affl-2* on HSF-1 subcellular localization and expression. While we did find some differences in HSF-1 expression prior to heat shock and HSF-1 granule formation after heat shock, we are not confident that these differences can explain the phenotypes of the various mutants because the distributions of our measurements for different genotypes overlap. This aligns with our hypothesis that *affl-2* is necessary for proper elongation in transcription, not initiation, for it suggests that AFFL-2 acts downstream of granule formation. Furthermore, addition of the FUS LC domain does not significantly increase the performance of the modified AFFL-2, which suggests that AFFL-2 is not involved in recruiting RNA polymerase but is acts in a downstream step. However, there are no putative binding sites for the granules and it is unclear what their role in heat shock response is (Morton and Lamitina 2013).

As mentioned previously, AFFL-1 and AFFL-2 are homologs of mammalian AFF1 and AFF4, which serve as scaffolds for the super elongation complex (SEC). AFF1 and AFF4 serve as scaffolds in the super elongation complex (SEC), which regulates release from promoter-proximal pausing during transcriptional elongation using P-TEFb (He and Zhou 2011; Lu *et al.* 2014; Mück *et al.* 2016). AFF4 is responsible for heat shock induced HSP70 expression, which illustrates that its role in heat shock induced gene expression is conserved (Lu *et al.* 2015). In *C. elegans,* the P-TEFb complex has been shown to be necessary for embryonic development and expression of *hsp-16.2* (Schulze-Gahmen *et al.* 2013). Although *affl-2* and *affl-1* mutants survive past embryonic development, *affl-2* mutants are deficient in heat shock induced gene expression. Future work should investigate whether *affl-2* mutants are also deficient in expression of genes involved in embryonic development and whether they have the same defects in Ser2 phosphorylation as animals lacking a function P-TEFb complex (Schulze-Gahmen *et al.* 2013).It is possible that *affl-2* mutants still have Ser2 phosphorylation, but at lower levels than wild type, which could allow them able to survive through development.

Our results demonstrate that the *C. elegans* ortholog of AF4/FMR2 family members, AFFL-2, is necessary for heat shock induced transcription. Our sequence analysis suggests that AF4/FMR2 homologues are found more widely in nature than previously thought, highlighting their importance. These results combined with previous work on other members of the SEC suggest that *C. elegans* can be a powerful, multicellular model to understand transcriptional elongation. Further study of *C. elegans* homologs of human AF4/FMR2 proteins will facilitate our understanding of heat shock response as well as transcriptional elongation in general.

## Acknowledgements

We thank Heenam Park for helping create *Y55B1BR.1* (*affl-1*) null mutants using CRISPR/Cas9, and we thank Jean Badroos, Jasmine S. Revanna, and Minyi Tan for technical assistance. We thank Hillel Schwartz, Jonathan Liu, and members of the Sternberg Lab for insightful discussions. We thank Sarah MacLean and the Sternberg Lab for comments on the manuscript. Some strains were provided by the CGC, which is funded by NIH Office of Research Infrastructure Programs (P40 OD010440). This work was supported by NIH (K99GM126137 to H.W., U24HG002223 to P.W.S., and R240D023041). The Millard and Muriel Jacobs Genetics and Genomics Laboratory at California Institute of Technology performed whole genome sequencing. S.J.W. was supported by the Caltech Student Faculty Programs, the family of Laurence J. Stuppy, Samuel N. Vodopia, and Carol J. Hasson.

## Data Availability

Strains and plasmids are available upon request. Code for data analysis and image processing can be found at: https://github.com/sophiejwalton/affl-2.git. Data will be made publicly available upon publication.

## Supporting Information

### Supplementary Data

Figure S1: *sup-45* (Y55B1BR.2) SNP mapping data.

Figure S2: Alignment of example AF4/FMR2 C-terminal Homology Domains.

Figure S3: Subcellular Localization of AFFL-2 N-terminal Deletion::GFP.

Figure S4: Morphology of AFFL-2 rescue variants.

Figure S5: Quantification of HSF-1::GFP in various mutants.

### Supplementary Experimental Procedures

**Appendix 1: Strains Used in This Study**

Table S1: Mutant Strains

Table S2: Strains used for *affl-2* expression experiments

Table S3: Strains used for *affl-2* rescue variants

Table S4: Strains for HSF-1::GFP localization experiments

**Appendix 2: Plasmids created for this study**

Table S5: pSJW003 Construction

Table S6: pSJW005 Construction

Table S7: pSJW0035 Construction

Table S8: pSJW0036 Construction

Table S9: pSJW0040 Construction

Table S10: pSJW0041 Construction

**Appendix 3: Oligos for genotyping used in this study**

