## Supplementary Information, Figs for "*Caenorhabditis elegans* AF4/FMR2 family homolog *affl-2* is required for heat shock induced gene expression"

**Supplementary Figures**

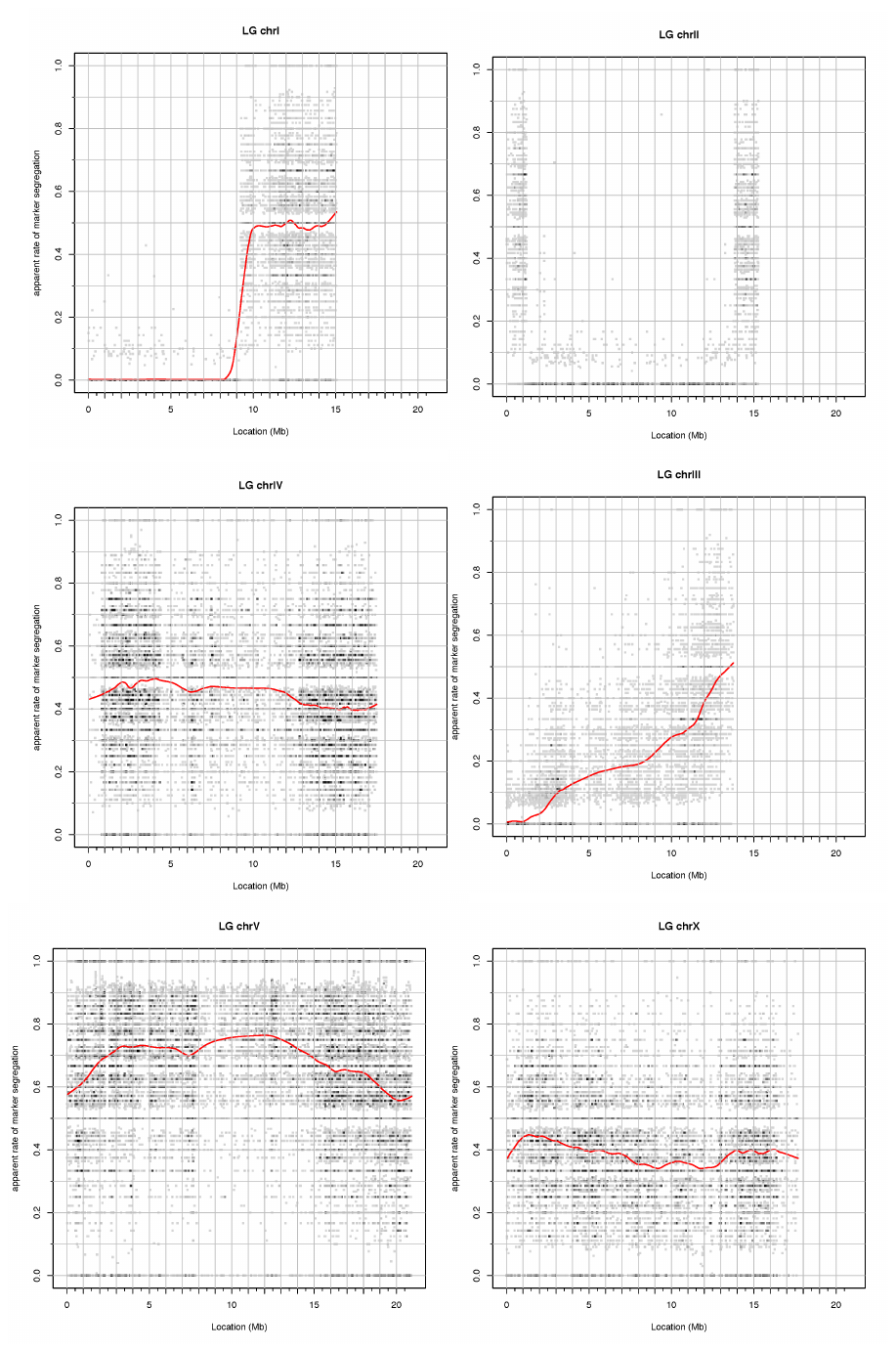

**Figure S1: *sup-45* (Y55B1BR.2) SNP mapping data.**

Mapping data for *sup-45(sy991*). The red line represents the rate of marker segregation. The rate is low in the left side of the chromosome I due to the *C. elegans* sperm toxicity gene *peel-1* (Seidel *et al.* 2011), and the rate is low in chromosome II due to the presence of *syIs231* that prevents recombination. Thus, the mutation is located in the left tip of chromosome III, where the rate is low as well.

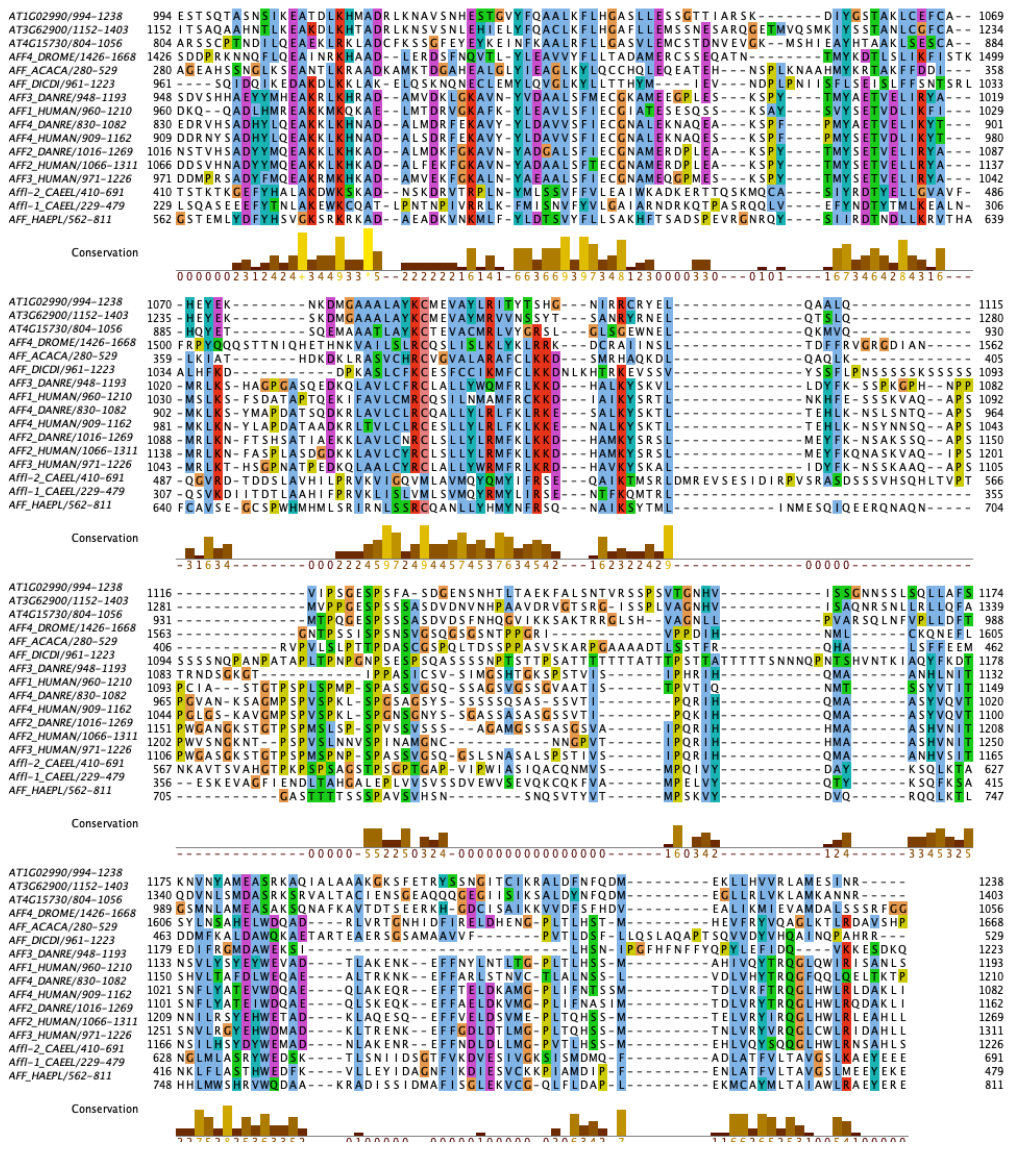

**Figure S2: Alignment of example AF4/FMR2 C-terminal Homology Domains.**

The multiple sequence alignment of CHD domains was generated using the MUSCLE software and rendered using the Jalview package with ClustalX colour scheme, where conserved residues are coloured as follows: hydrophobic (blue), positive charge (red), negative charge (magenta), polar (green), cysteines (pink), glycine (orange), proline (yellow) and aromatic (cyan). The UniProtKB accession identifiers for each sequence are listed here: At1g02990, F4HZA0; At3g62900, F4IZK5; At4g15730, Q8GY51; AFF4_DROME, Q9VQI9; AFF_ACACA, L8H858; AFF_DICDI, Q54PM6; AFF3_DANRE, F1RA06; AFF1_HUMAN, P51825; AFF4_DANRE, I3ISK1; AFF4_HUMAN, Q9UHB7; AFF2_DANRE, E7F2E1; AFF2_HUMAN, P51816; AFF3_HUMAN, P51826; Affl-2_CAEEL, Q95XW7; Affl-1_CAEEL, Q95XW6; AFF_HAEPL, A0A158QQA2.

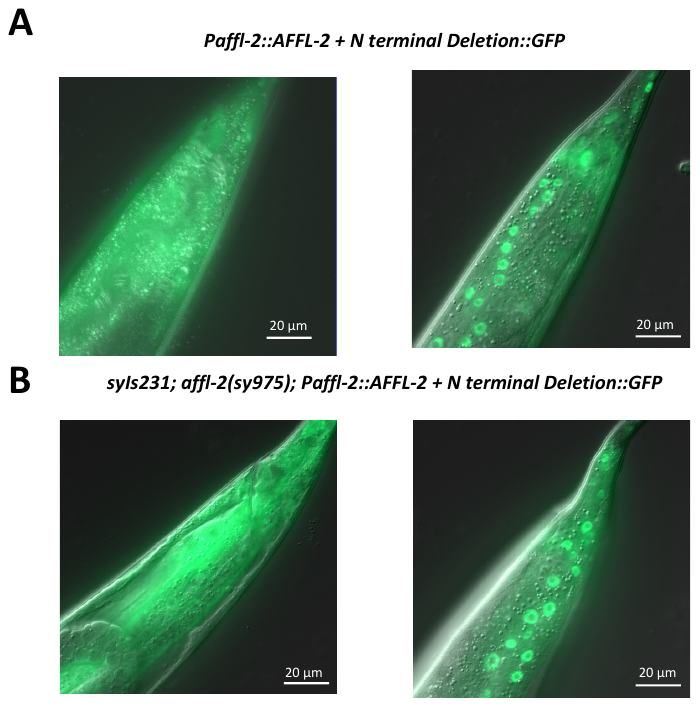

**Figure S3: Subcellular Localization of AFFL-2 N-terminal Deletion::GFP.**

Localization of AFFL-2 N-terminal Deletion::GFP in wild type (A) and in *affl-2(sy975)* mutants (B). All photos were taken of young adult worms at room temperature.

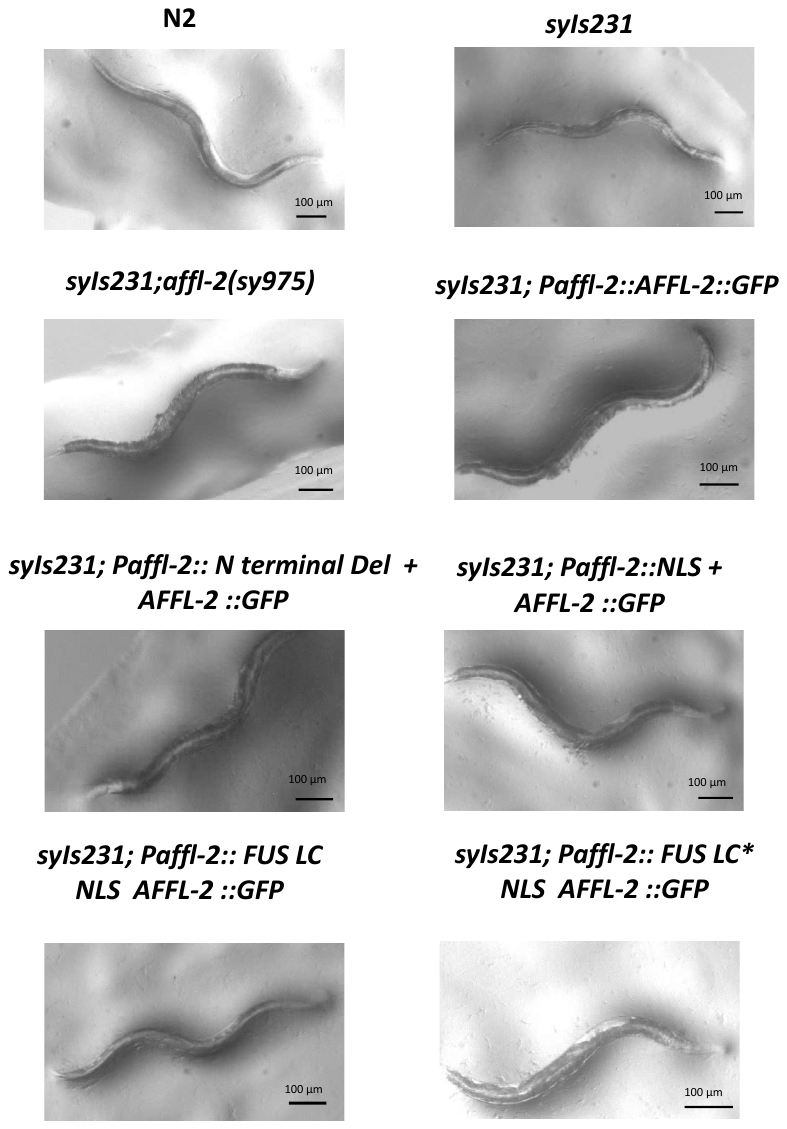

**Figure S4: Morphology of AFFL-2 rescue variants.**

Images of wild type (N2), *syIs231, syIs231; sup-45(sy975),* and *syIs231; sup-45(sy975)* animals with indicated rescue constructs. All constructs are arrays.

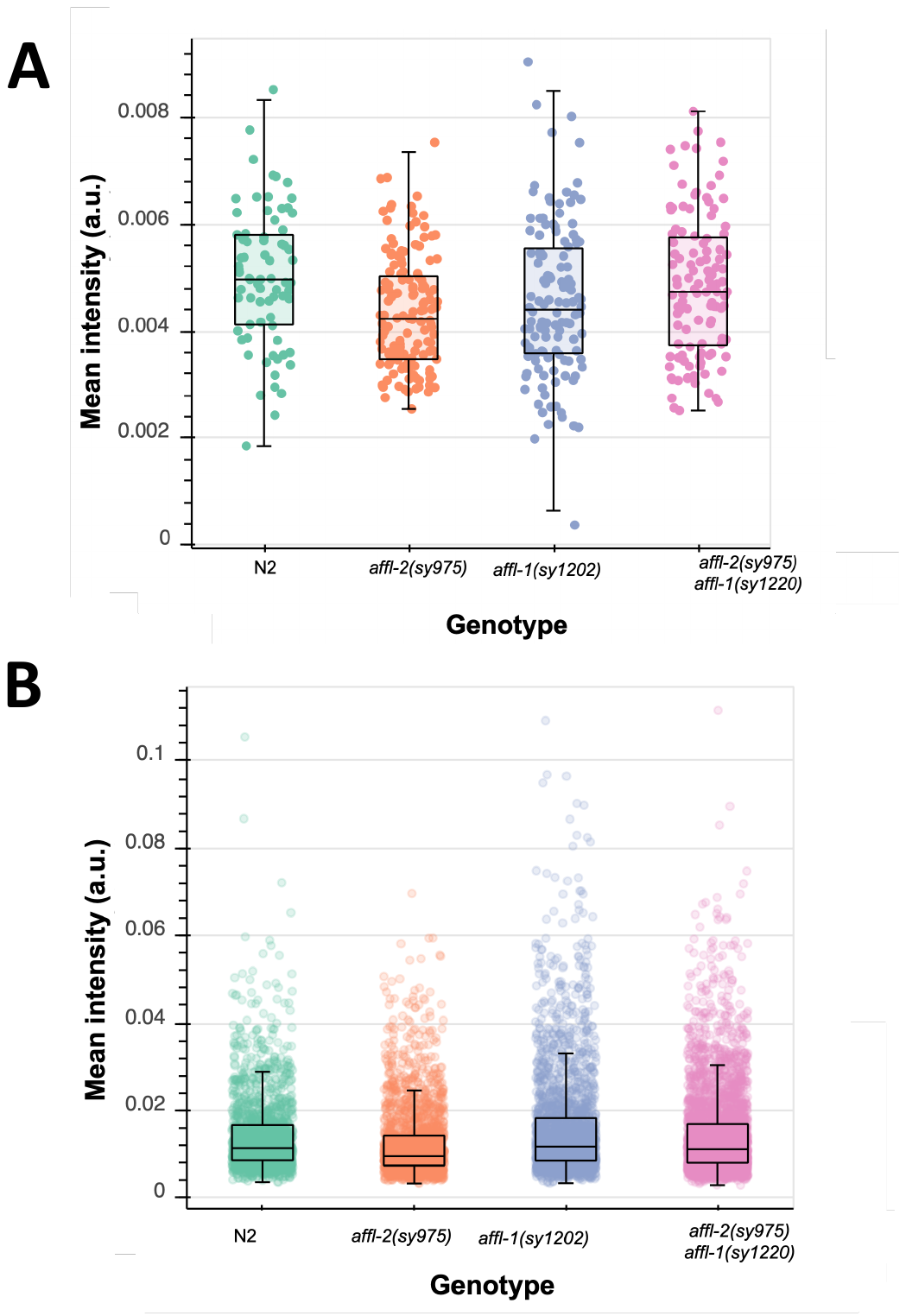

**Figure S5: Quantification of HSF-1::GFP in various mutants.** A) Mean intensity (a.u.) of HSF-1::GFP in nuclei in animals at 20 °C. B) Mean intensity (a.u.) of HSF-1::GFP nuclear granules formed after a five-minute heat shock at 35°C.

**Supplementary Experimental Procedures**

**Appendix 1: Strains Used in This Study**

The wild type strain was N2 Bristol and the parental strain for EMS screen wasPS7244 (*syIs231*).

Our mapping strain was PS7421, which was constructed by outcrossing PS7244 10 times to the Hawaiian mapping strain CB4856, so that *syIs231* is in a Hawaiian background.

**Table S1: Mutant Strains**

| Name | Genotype | Comments/Citation |
| --- | --- | --- |
| PS3551 | *hsf-1(sy441)I* | Hajdu-Cronin *et al.* 2004 |
| PS8572 | *hsf-1(sy441)I; syIs231 II* | Constructed by crossing PS3551 and PS7244 |
| PS7497 | *hsf-1(sy1198) I; syIs231 II* | Isolated in this screen |
| PS3233 | *affl-2(sy509) III* | Hajdu-Cronin *et al.* 2004 |
| PS8082: | *syIs231 II; affl-2 (sy975) III* | Isolated in this screen: outcrossed 4x. |
| PS8060 | *affl-2(sy975) III* | Isolated in this screen: outcrossed 4x. |
| PS7594 | *syIs231 II; affl-2(sy981) III* | Isolated in this screen |
| PS7644 | *syIs231 II; affl-2(sy991) III; syIs197 V* | Isolated in this screen |
| PS2323 | *affl-2(sy439) III; dpy-20(e1282) syIs17 IV* | Hajdu-Cronin *et al.* 2004 |
| PS8176 | *affl-1(sy1202)* | STOP-IN cassette (Wang *et al.* 2018) knocked into *affl-1* in N2 |
| PS8203 | *affl-2(sy975) affl-1(sy1220) III* | STOP-IN cassette (Wang *et al.* 2018) knocked into *affl-1* in PS8060. |

***affl-2* driver:**

*affl-2* driver strains were constructed using strains and methods from (Wang *et al.* 2017). To create *syEx1666*, 25 ng/ul of pSJW003 along with 40 ng/ µl of P*unc-122::rfp* and 35 ng/µl of 1 kb DNA ladder (NEB) was injected into *syIs300*.

**Table S2: Strains used for *affl-2* expression experiments**

| Name | Genotype | Comments/Citation |
| --- | --- | --- |
| **PS7965** | *syEx1666 [pSJW003(Paffl-2::nls::GAL4SK::VP64::let-858 3'UTR), 25ng/µL; Coel::RFP, 40ng/µL; 1kb DNA ladder (NEB), 35 ng/µL]; ]; syIs300[pG4US7(15xUAS:: pes-10::gfp::unc-54 3'UTR), 25ng/µL; Pttx-3::rfp, 40ng/µL; pBlueScript, 35ng/µL] V* |  |
| **PS8573** | *syEx1666 [pHW494(15xUAS::Δpes-10::gfp::H2B::let-858 3'UTR), 25ng/µL; Pttx-3::rfp, 40ng/µL; pBlueScript, 35ng/µL]; syIs407* | Constructed by outcrossing PS7957 to remove syIs300, and then crossing *syEx1666* animals with the GFP::H2B effector strain PS7186 (*syIs407*). |

***affl-2* rescue variants:** All strains were constructed by injecting 10 ng/µL of plasmid containing the rescue construct along with 80 ng/µl of plasmid bluescript and 10 ng/ µL of the coninjection marker KP1368 into PS8082 (*syIs231 II; affl-2 (sy975) III*), and then outcrossing to remove PS8082 if indicated.

**Table S3: Strains used for *affl-2* rescue variants**

| Name | Genotype | Comments/Citation |
| --- | --- | --- |
| PS8122 | *syIs231 II; affl-2 (sy975) III*; *syEx1668*  *[Paffl-2::affl-2 cDNA::GFP::unc-54 3'UTR); KP1368,; pBluescript]* | Plasmid containing rescue construct is pSJW005. |
| PS8137 | *syEx1668* | Constructed by outcrossing PS8122 |
| PS8242 | *syIs231 II; affl-2 (sy975) III*; *syEx1733-*  *[Paffl-2::N term deletion affl-2 cDNA::GFP::unc-54 3'UTR); KP1368; pBluescript]* | Plasmid containing rescue construct is pSJW0035. |
| PS8275 | *syEx1733* | Constructed by outcrossing PS8242 |
| PS8243 | *syIs231 II; affl-2 (sy975) III*; *syEx1734*  *[Paffl-2::N term deletion + NLS affl-2 cDNA::GFP::unc-54 3'UTR); KP1368; pBluescript]* | Plasmid containing rescue construct is pSJW0036. |
| PS8276 | *syEx1734* | Constructed by outcrossing PS8243 |
| PS8325 | *syIs231 II; affl-2 (sy975) III*; *syEx1727[pSJW0040(Paffl-2::FUS NLS sup- 45 cDNA::GFP::unc-54 3' UTR, 10 ng/ L; KP1368, 10ng/ L; pBluescript, 80 ng/ L]* | Plasmid containing rescue construct is pSJW0040. |
| PS8491 | *syEx1727* | Constructed by outcrossing PS8325 |
| PS8492 | *syIs231 II; affl-2 (sy975) III*; *syEx1735*  *[Paffl-2:: N term deletion + FUS Variant + NLS + affl-2 cDNA::GFP::unc-54 3'UTR); KP1368; pBluescript]* | Plasmid containing rescue construct is pSJW0041. |
| PS8493 | *syEx1735* | Constructed by outcrossing PS8492 |

**HSF-1::GFP localization experiments:** All strains with *drSi13* (except OG497), where constructed through crossing the indicated strain with OG497 using BN578 *bqSi189 [lmn-1p::mCherry::his-58 + unc-119(+)]; bqSi577 [myo-2p::GFP + unc-119(+)]* as a balancer. Genotypes were verified using PCR.

**Table S4: Strains for HSF-1::GFP localization experiments**

| Name | Genotype | Comments/Citation |
| --- | --- | --- |
| OG497 | *drSi13 II [phsf-1::hsf-1::GFP::unc-54 3'UTR + Cbr-unc-119(+)]; unc-119(ed3) III* | Morton and Lamitina 2013 |
| PS8272 | *drSi13 II; affl-2(sy975)** | Constructed by crossing PS8082 and OG497 |
| PS8273 | *drSi13 II; affl-1(sy1202)** | Constructed by crossing PS8176 and OG497 |
| PS8274 | *drSi13 II; affl-2(sy975); affl-1(sy1220); unc-119(ed3)** | Constructed by crossing PS8203 and OG497 |

***:** drSi13 rescues unc-119(ed3) phenotype. We are not sure if these strains still have the unc-119(ed3) after outcrossing.

**Appendix 2: Plasmids created for this study**

pSJW003: P*affl-2*::cGAL::let-858 3'UTR

Created by T4 Ligation with P*affl-2* and the vector pHW393 (Wang *et al.* 2017)

Both the vector and the insert were digested with FseI and AscI.

**Table S5: pSJW003 Construction**

| Fragment | Primers | Comments/Citation |
| --- | --- | --- |
| P*affl-2* | oSJW33f:  cccGGCCGGCCcggagctattgattcaggattg | Template is N2 genomic DNA. |
|  | oSJW34r:  cccGGCGCGCCTTCCCATtgaaatcgtcttcccg |  |

pSJW005: P*affl-2*::*affl-2* cDNA::GFP::*unc-54 3'UTR*

Created by Gibson Assembly with the vector pHW447 (digested with FseI and NotI) and P*affl-2*::*affl-2* cDNA fragment.

**Table S6: pSJW005 Construction**

| Fragment | Primers | Comments/Citation |
| --- | --- | --- |
| P*affl-2*::*affl-2* cDNA | oSJW82f:  GAAATGAAATAAGCTTGCATGCGGCCGGCCcggagct  attgattc | Template is pSJW004 |
|  | oSJW83r:  GTTCTTCTCCTTTACTACTtGCGGCCGCTTGCTTCC  GACGAGCTTTTC |  |

pSJW035: P*affl-2*::N term deletion *affl-2* cDNA::*GFP*::*unc-54 3'UTR*

Created by Gibson Assembly with pSJW005 digested with PacI and XbaI. Two Inserts were obtained via PCR from pSJW005 in order to create a deletion of both NLSs predicted by cNLS software (Kosugi *et al.* 2009).

**Table S7: pSJW0035 Construction**

| Fragment | Primers | Comments/Citation |
| --- | --- | --- |
| P*affl-2*:: *affl-2* cDNA 5’ end | oSJW163F: ggcaactttgacctatcttaattaatttttaaactataattcgataatttttcgag | Template is pSJW005 |
|  | oSJW162R:  TGAGTCTTCTTCCATtaaatatg |  |
| *affl-2* cDNA 3’ end | oSJW165F:  catatttaATGGAAGAAGACTCAAGGAATTACATTCCCCCTTC | Template is pSJW005 |
|  | oSJW164R:  CCTGGGCCAAGTATTGGGTCTCTAGAAGGTGTGGGTACCGCAG |  |

**Table S8: pSJW0036 Construction**

pSJW036: P*affl-2*:: NLS + *affl-2* cDNA::*GFP::unc-54 3'UTR*

Created by Gibson Assembly with pSJW005 digested with PacI and XbaI. Two Inserts were obtained via PCR from pSJW005 in order to substitute an artificial NLS (Wang *et al.* 2017) for a deletion of both NLSs predicted by cNLS software (Kosugi *et al.* 2009).

| Fragment | Primers | Comments/Citation |
| --- | --- | --- |
| P*affl-2*:: *affl-2* cDNA 5’ end | oSJW163F: ggcaactttgacctatcttaattaatttttaaactataattcgataatttttcgag | Template is pSJW005 |
|  | oSJW166R:  CCTTTTACGCTTCTTTTTAGGTGAGTCTTCTTCCATtaaatatg |  |
| NLS + *affl-2* cDNA 3’ end | oSJW167F:  CTCACCTAAAAAGAAGCGTAAAAGGAATTACATTCCCCCTTC | Template is pSJW005 |
|  | oSJW164R:  CCTGGGCCAAGTATTGGGTCTCTAGAAGGTGTGGGTACCGCAG |  |

pSJW040: P*affl-2*:: FUS LC + NLS *+ affl-2* cDNA::*GFP::unc-54 3'UTR*

Created by Gibson Assembly with pSJW005 digested with PacI and XbaI. Two Inserts were obtained via PCR from pSJW005 in order to substitute an artificial NLS for a deletion of both NLSs predicted by cNLS software (Kosugi *et al.* 2009; Wang *et al.* 2017). In between the two inserts, the FUS LC domain was inserted (Kwon *et al.* 2013).

**Table S9: pSJW0040 Construction**

| Fragment | Primers | Comments/Citation |
| --- | --- | --- |
| P*affl-2*:: *affl-2* cDNA 5’ end | oSJW163F: ggcaactttgacctatcttaattaatttttaaactataattcgataatttttcgag | Template is pSJW005 |
|  | oSJW162R:  TGAGTCTTCTTCCATtaaatatg |  |
| FUS LC domain | oSJW169F: catatttaATGGAAGAAGACTCAgcctcaaacgattatacccaac | Domain from Kwon *et al.* 2013 |
|  | oSJW174R: CCTTTTACGCTTCTTTTTAGGcccacggtcctgctgtccatag |  |
| NLS + *affl-2* cDNA 3’ end | oSJW173F: gggCCTAAAAAGAAGCGTAAAAGGAATTACATTCCCCCTTC | Template is pSJW005 |
|  | oSJW164R:  CCTGGGCCAAGTATTGGGTCTCTAGAAGGTGTGGGTACCGCAG |  |

pSJW041: P*affl-2*:: FUS LC * + NLS + *affl-2* cDNA::*GFP::unc-54 3'UTR*

Created by Gibson Assembly with pSJW005 digested with PacI and XbaI. Two Inserts were obtained via PCR from pSJW005 in order to substitute an artificial NLS for a deletion of both NLSs predicted by cNLS software (Kosugi *et al.* 2009; Wang *et al.* 2017)

In between the two inserts, the modified FUS LC domain (FUS LC *) was inserted (Kwon *et al.* 2013).

**Table S10: pSJW0041 Construction**

| Fragment | Primers | Comments/Citation |
| --- | --- | --- |
| P*affl-2*:: *affl-2* cDNA 5’ end | oSJW163F: ggcaactttgacctatcttaattaatttttaaactataattcgataatttttcgag | Template is pSJW005 |
|  | oSJW162R:  TGAGTCTTCTTCCATtaaatatg |  |
| FUS LC * | oSJW169F: catatttaATGGAAGAAGACTCAgcctcaaacgattatacccaac | Domain from Kwon *et al.* 2013 |
|  | oSJW174R: CCTTTTACGCTTCTTTTTAGGcccacggtcctgctgtccatag |  |
| NLS + *affl-2* cDNA 3’ end | oSJW173F: gggCCTAAAAAGAAGCGTAAAAGGAATTACATTCCCCCTTC | Template is pSJW005 |
|  | oSJW164R:  CCTGGGCCAAGTATTGGGTCTCTAGAAGGTGTGGGTACCGCAG |  |

**Appendix 3: Oligos for genotyping used in this study**

All primers are listed from 5’ to 3’.

***affl-2*:**

First, the following external primers were used to PCR out sections of *affl-2*:

Fragment 1:

oSJW26F: TTTACGGCATTTGGGCTTTGG

oSJW45R: GGCGGAGCTTGTAACGTGAC

Fragment 2:

oSJW12F: GGTGGAGAGACGCAGAGTTC

oSJW14R: TAATCCTAAGGCAAAGCCCA

Fragment 3:

oSJW26F: TTTACGGCATTTGGGCTTTGG

oSJW15R: CAGTGTGACTAGAGAGGAGAAC

The following internal primers were used for Sanger Sequencing:

oSJW47R: TCGACTATTTGCCGGTTTGC

oSJW12F: GGTGGAGAGACGCAGAGTTC

oSJW41R: TCCCACTACGGTTTGATCTAC

oSJW46F: TCAAAGTGGTGTGCAAGGAAGATG

oSJW75R: ACTGGAACCTTGAAGCCCTTG

oSJW13R: CCAAAGCCTAAGCCTGAGC

oSJW42F: TCGGCTCGCTTTAGGTTTATC

oSJW152R: CTAGCATCACCTGACCGATCAC

oSJW43F: GCCCAGGGACTAAGACTAAAC

oSJW44R: GTAGGCATCGTAGACAATTTGG

oSJW15R: CAGTGTGACTAGAGAGGAGAAC

***hsf-1*:**

First, the following external primers were used to PCR out sections of *hsf-1*:

Fragment 1:

oSJW35F: TGCCGGTATTGCCGAATTTG

oSJW62R: TCGTCGTCAACTTTGTTGTTTC

Fragment 2:

oSJW63F: AAACAACAAAGTTGACGACGAC

oSJW65R: TATTGCTGTTGGCGAGCATG

Fragment 3:

oSJW64F: AGTAATGGCAGAGATGCGTG

oSJW37R: GATGGAGCCGAAGATGATGT

Fragment 4:

oSJW40F: AACGTGCTCGAATGAACTCTG

oSJW67R: TCAAGAGCAAGCTGTCTGAG

Fragment 5:

oSJW66F: TCTATATTCTCCAACTCTCGGACTCTC

oSJW39R: CCTCTCCCATCTTCCATCTCCATC

The following internal primers were used for Sanger Sequencing:

oSJW48F: TTACCTCTATTGCCGAGTTTGC

oSJW70F: TTAGGGAAAGTGACGGAACACG

oSJW79R: ACATATTCAACTGTCTGACCATGC

oSJW52F: ATTTTTACACGGAGATACCC

oSJW53R: GCCATTTCGAGAAATTTGCG

oSJW81R: TCCTGAGCACTTCCTCTTCCAAC

oSJW37R: GATGGAGCCGAAGATGATGT

oSJW61F: TGTTGGAAGAGGAAGTGCTCAGG

oSJW50F: GCAAGCTCCGCCCATTTATTG

oSJW67R: TCAAGAGCAAGCTGTCTGAG

oSJW66F: TCTATATTCTCCAACTCTCGGACTCTC

oSJW80F: TGGAACTGATACTTCATTGGAGAG

oSJW39R: CCTCTCCCATCTTCCATCTCCATC

***affl-1*:**

Genotyping methods were adapted from (Wang *et al.* 2018).

Briefly, oHP083F and oHP013R will only produce product if the STOP-IN cassette is present.

oHP083F and oHP084R will produce product regardless, but of different lengths depending on the presence of the STOP-IN cassette.

Fragment 1:

oHP083F: AACTCAAATTCCAGGGCAAACC

oHP013R: GCTTATCACTTAGTCACCTCTGCTC

Fragment 2:

oHP083F: AACTCAAATTCCAGGGCAAACC

oHP084R: TGGTGAGGTAGCCGTTGAATC

**Supplementary Section References**

Hajdu-Cronin Y. M., W. J. Chen, and P. W. Sternberg, 2004 The L-type cyclin CYL-1 and the heat-shock-factor HSF-1 are required for heat-shock-induced protein expression in Caenorhabditis elegans. Genetics 168: 1937–1949. https://doi.org/10.1534/genetics.104.028423

Kosugi S., M. Hasebe, M. Tomita, and H. Yanagawa, 2009 Systematic identification of cell cycle-dependent yeast nucleocytoplasmic shuttling proteins by prediction of composite motifs. Proc. Natl. Acad. Sci. U.S.A. 106: 10171–10176. https://doi.org/10.1073/pnas.0900604106

Kwon I., M. Kato, S. Xiang, L. Wu, P. Theodoropoulos, *et al.*, 2013 Phosphorylation-Regulated Binding of RNA Polymerase II to Fibrous Polymers of Low-Complexity Domains. Cell 155: 1049–1060. https://doi.org/10.1016/j.cell.2013.10.033

Morton E. A., and T. Lamitina, 2013 Caenorhabditis elegans HSF-1 is an essential nuclear protein that forms stress granule-like structures following heat shock. Aging Cell 12: 112–120. https://doi.org/10.1111/acel.12024

Seidel H. S., M. Ailion, J. Li, A. van Oudenaarden, M. V. Rockman, *et al.*, 2011 A Novel Sperm-Delivered Toxin Causes Late-Stage Embryo Lethality and Transmission Ratio Distortion in C. elegans. PLOS Biology 9: e1001115. https://doi.org/10.1371/journal.pbio.1001115

Wang H., J. Liu, S. Gharib, C. M. Chai, E. M. Schwarz, *et al.*, 2017 cGAL, a temperature-robust GAL4–UAS system for *Caenorhabditis elegans*. Nature Methods 14: 145–148. https://doi.org/10.1038/nmeth.4109

Wang H., H. Park, J. Liu, and P. W. Sternberg, 2018 An Efficient Genome Editing Strategy To Generate Putative Null Mutants in Caenorhabditis elegans Using CRISPR/Cas9. G3: Genes, Genomes, Genetics 8: 3607–3616. https://doi.org/10.1534/g3.118.200662
